# Altered physiological, affective, and functional connectivity responses to acute stress in patients with alcohol use disorder

**DOI:** 10.1101/2024.01.18.576207

**Authors:** Yana Schwarze, Johanna Voges, Alexander Schröder, Sven Dreeßen, Oliver Voß, Sören Krach, Frieder Michel Paulus, Klaus Junghanns, Lena Rademacher

## Abstract

**Background:** There is evidence that the processing of acute stress is altered in alcohol use disorder (AUD), but little is known about how this is manifested simultaneously across different stress parameters and which neural processes are involved. The present study examined physiological and affective responses to stress and functional connectivity in AUD.

**Methods:** Salivary cortisol samples, pulse rate, and affect ratings were collected on two days from 34 individuals with moderate or severe AUD and 34 controls. On one day, stress was induced, and on the other day, a non-stressful control task was performed. Following the intervention, participants underwent fMRI to assess functional connectivity, focusing on cortical and subcortical seed regions previously reported to be involved in AUD and/or stress.

**Results:** For pulse rate and cortisol, stress responses were blunted in AUD, whereas negative affect was increased. Furthermore, stress-related changes in pulse rate, cortisol, and affect were only correlated in healthy controls. Neuroimaging analyses revealed stress-related group differences in functional connectivity, involving the connectivity of striatal seeds with the posterior DMN, cerebellum and midcingulate cortex, and of the posterior DMN seed with the striatum and thalamus.

**Conclusions:** The results suggest a dissociation between subjective experienced distress and the physiological stress response in AUD as well as stress-related alterations in functional connectivity. These findings highlight the complex interplay between chronic alcohol use and acute stress regulation, offering valuable considerations for the development of therapeutic strategies.

## Introduction

There is a complex and bidirectional relationship between stress and alcohol use disorder (AUD). On the one hand, stress is a known risk factor for the development of addiction and the vulnerability to relapse (1). On the other hand, chronic alcohol consumption is associated with alterations of physiological stress systems, including the hypothalamic-pituitary-adrenal (HPA) axis and the autonomic nervous system: Basal cortisol levels in blood plasma and saliva, which serve as indicators of HPA axis activity, have been observed to be elevated in patients diagnosed with AUD during intoxication and acute withdrawal (e.g. (2,3)). During abstinence, cortisol levels generally decline (4,5) and most studies do not find significant group differences anymore (6). Regarding the autonomic nervous system (ANS), decreased heart rate variability and increased heart or pulse rate have been reported for early abstinence (7–11).

In addition to these alterations in basal ANS and HPA axis tone, phasic responses to stress have also been found to be disturbed: According to recent reviews (6,12), numerous studies indicate that the cortisol response to acute stress is blunted in AUD, impacting relapse risk (13–15). Furthermore, a decreased heart (or pulse) rate response to acute stress has been observed in several studies, although the results here are more mixed (6). Interestingly, a diminished physiological stress response does not appear to be associated with lower subjective perceived stress. Few studies have examined effects of acute stress on subjective distress or affect, but these report equal or increased negative affect or distress in AUD compared to healthy controls (7,10,16–18). Thus, previous findings suggest a dissociation between physiological and subjective stress responses in AUD: while there is evidence that HPA and ANS responses are decreased, experienced distress tends to be higher than in healthy subjects.

So far, it is not clear which neural processes underlie altered stress responses. The investigation of functional connectivity using functional magnetic resonance imaging (fMRI) can provide valuable insights into the organization of brain circuits (19). To our knowledge, there has only been one pilot study of functional connectivity during acute stress in ten long-term abstinent patients with AUD and eleven controls (20). This study focused on the amygdala as seed region and found decreased connectivity with frontal, temporal, parietal, and cerebellar regions in AUD while performing a classic stress task. However, other seed regions may also be of interest such as the thalamus and posterior cingulate cortex (PCC) - a central node in the default mode network (DMN) - where changes in connectivity have been demonstrated after stress induction in healthy subjects (for review see (21)). Furthermore, Wade and colleagues did not contrast the stress task with a non-stressful control session so it remains unclear to what extent the changes found are specific to a state of acute stress. Several other studies have also identified differences in functional connectivity between individuals with AUD and healthy controls in the absence of stress, including hyperconnectivity of the striatum with the anterior insula (22), anterior cingulate cortex (ACC), and superior and inferior frontal gyri (22,23), hypoconnectivity of the thalamus with the striatum (22–24), medial prefrontal cortex (24,25), and ACC (23,24,26), as well as disrupted connectivity of the DMN (27).

The present study addresses the gap in research on stress responses in AUD by investigating physiological and affective responses along with brain connectivity. Patients with moderate or severe AUD and healthy controls were examined on two separate days with a stress-inducing task on one day and a non-stressful control task on the other (counterbalanced in order). Salivary cortisol samples, pulse rate, and questionnaire data on negative affect were collected. Participants underwent fMRI to assess stress-induced changes in functional connectivity, focusing on cortical and subcortical regions that have been associated with AUD. We expected to find a dissociation between physiological and subjective stress responses in AUD which is reflected in increased affective, but blunted cortisol and pulse rate responses compared to healthy controls. Furthermore, we tested group differences in stress-associated changes in functional connectivity.

## Methods and Materials

### Sample

Forty-two individuals with a DSM-V diagnosis of moderate or severe alcohol use disorder (5 to 11 criteria fulfilled) and 38 healthy controls were recruited between February 2019 and October 2022 with a pause between March 2020 and October 2021 due to the SARS-CoV-19 pandemic (see Supplement for details on recruitment and inclusion and exclusion criteria). All patients were in the phase of early abstinence (10th - 40th day) after acute withdrawal symptoms had resolved and remained stationary in the clinic throughout the study. A final sample of 34 patients and 34 healthy controls was included in the study (see Supplements for details on dropouts and exclusions). The two groups were matched regarding age and gender. All participants gave informed and written consent. The study was approved by the local ethics committee at the University of Lübeck (AZ 17-077).

### Procedure

Before participating in the study, participants were informed about the study procedure (but not about the stress induction protocol) and screened to ensure they met all inclusion and exclusion criteria. After successful inclusion, participants visited the Center of Brain, Behavior and Metabolism in Lübeck on two separate days within a maximum interval of 10 days (mean interval: 3.25 days) for all but one person (see Supplement for details).

The two test days were identical in procedure and differed only in the stress protocol, with the order of the two days being counterbalanced across subjects (Figure 1A): On one day, participants completed the Trier Social Stress Test (TSST; (28)) as stress induction paradigm, on the other day a control task without stress induction (for details on the tasks see Supplement). Each test day lasted approximately three hours (8:30-11:30 am +/- 30 minutes) and started with a check of the inclusion and exclusion criteria and the collection of a first saliva sample (T0)^1^. This was followed by an explanation and practice of the experimental tasks to be performed later during the MRI session. Afterwards, a pulse oximeter (PULOX PO-300, Novidion GmbH) was attached to the participant’s index finger to record the pulse rate and the participants watched a calming video showing landscape scenes. Following this rest period, a second saliva sample was collected (T1) and participants completed the state part of the State-Trait-Anxiety Inventory (STAI-S, (29)) to assess their current affective state (see Supplements). Then the TSST or control task was performed. Another saliva sample (T2) and affective state rating were then collected and the pulse oximeter was removed. Another saliva sample (T3) was taken right before the start of the one-hour MRI session. Finally, participants provided a final saliva sample (T4), completed several questionnaires (see Supplements), and were debriefed on the day of the stress induction. On the second testing day, participants received their remuneration of €60 for their participation in the study.

**Figure 1.**
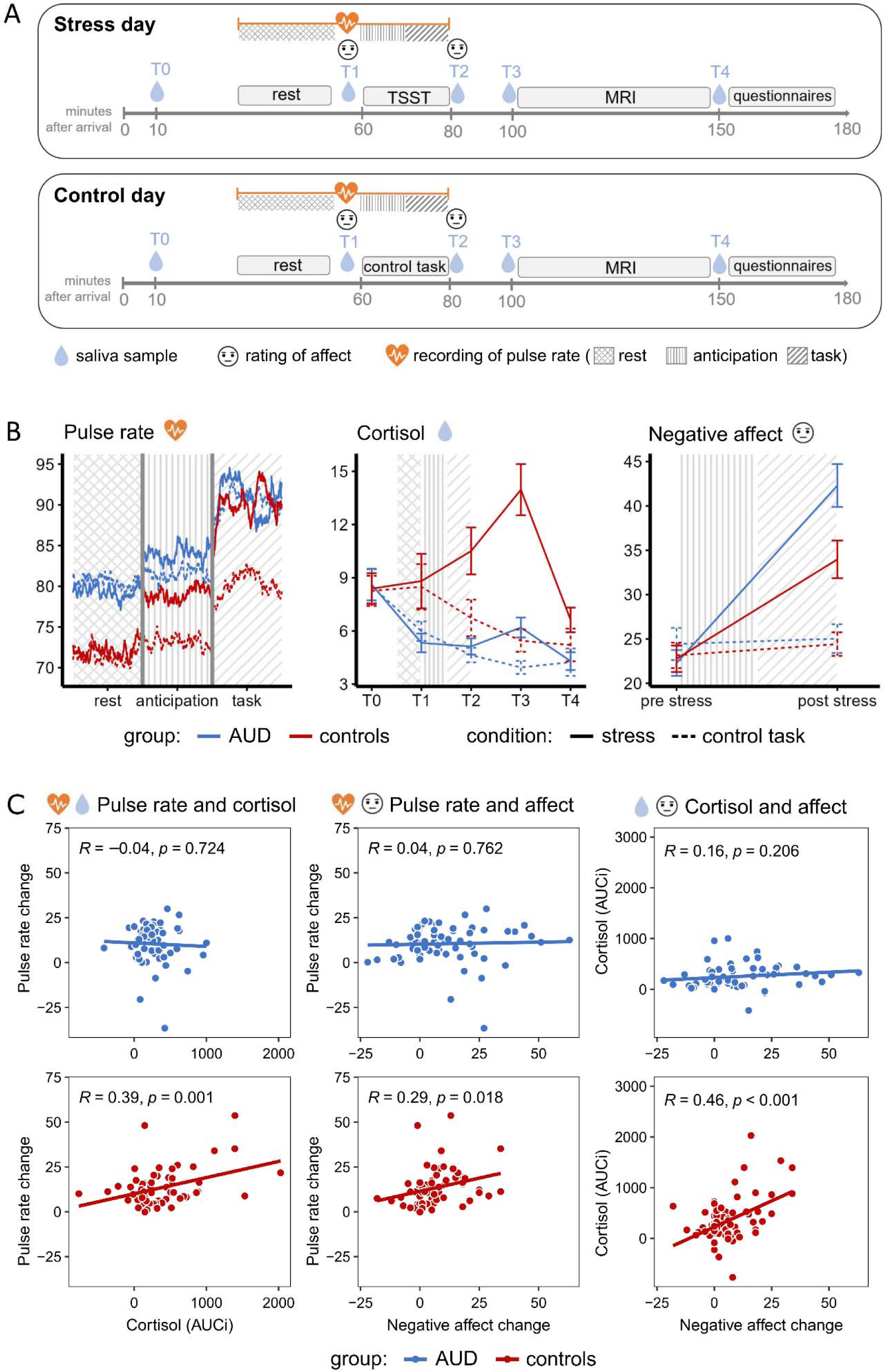
(A) Overview of the experimental design. (B) Development of pulse rate (bpm), cortisol levels (nmol/l), and rated negative affect over the two test days in alcohol use disorder (AUD) and controls. The shaded areas indicate the time of the rest phase as well as the anticipation and task phase of the stress or control task. The ANOVA on difference scores of pulse rate (task phase - rest) yielded a significant main effect of *condition* (*F*(1, 64) = 13.75, *p* < .001, η*_p_^2^* = .18) and a significant interaction effect of *condition* and *group* (*F*(1, 64) = 7.43, *p* = .008, η*_p_^2^* = .10), but no significant main effect of *group* (*F*(1, 64) = 1.47, *p* = .230, η*_p_^2^* = .02). The ANOVA on changes in cortisol levels (AUCi) also revealed a significant main effect of *condition* (*F*(1, 66) = 30.35, *p* < .001, η*_p_^2^* = .32) and an interaction effect of *condition* and *group* (*F*(1, 66) = 6.62, *p* = .012, η*_p_^2^*= .09), but no main effect of *group* (*F*(1, 66) = 2.79, *p* = .099, η*_p_^2^* = .04). The ANOVA on self-reported changes in affect (STAI-S after the task - STAI-S before the task) again revealed a significant main effect of *condition* (*F*(1, 66) = 72.12, *p* < .001, η*_p_^2^* = .52) and a significant interaction effect of *condition* and *group* (*F*(1, 66) = 7.50, *p* = .008, η*_p_^2^*= .10), but no main effect of *group* (*F*(1, 66) = 3.73, *p* = .058, η*_p_* = .05). (C) Relationship of stress-induced changes in cortisol, pulse rate, and negative affect. The three parameters correlate significantly in the control group, but in AUD there is no association. Comparisons of correlations revealed a significantly stronger association of stress-related changes in cortisol and pulse rate (*p* = 0.009) and of cortisol and negative affect (*p* = 0.049) in healthy subjects than in AUD. The comparison of the correlation of pulse rate and negative affect did not reach significance (*p* = 0.138).

Due to the SARS-CoV-19 pandemic, data collection was temporarily paused between March 2020 and October 2021 and some procedural changes were made after the pause (see Supplement for details). We reran all analyses with a covariate distinguishing between measurements taken before and during the pandemic, which yielded no substantial changes in the results (see Supplement).

### Analysis of pulse rate, cortisol, and affective state

For every participant and test day, mean scores of the STAI-S were calculated for the two time points. Mean pulse rate scores for the resting, the anticipation, and the task phase were also computed (see Supplements). Two control subjects had to be excluded from analyses involving pulse rate due to incomplete recording. Cortisol concentrations for saliva samples were measured by chemiluminescence immunoassay with high sensitivity (see Supplements for details on laboratory analysis).

To assess group differences in basal activity of the sympathetic nervous system, HPA activity, and subjectively experienced distress, averaged baseline scores for cortisol (T0), pulse rate, and STAI-S of the two test days were compared between the groups using two-tailed Welch’s two-sample t-tests.

To analyze the effects of the experimental stress induction, repeated measures analyses of variance (ANOVA) with *group* (AUD vs. controls) as the between-subject and *condition* (stress vs. control task) as the within-subject factor were calculated on difference scores of pulse rate (mean pulse rate during task phase - mean pulse rate during rest phase), difference scores of self-reported affective state (STAI-S after stress induction/control task - STAI-S before the task), and changes in cortisol (area under the curve with respect to increase (AUCi) from T1 to T4 (for reference see (30))).

To explore the association between stress-related changes in pulse rate, cortisol, and negative affect, Pearson’s correlations were computed for the three difference scores described above and correlations within the two groups (AUD and controls) were statistically compared (31).

### MRI acquisition and data processing

Participants were examined in the MRI at the Center for Brain, Behavior and Metabolism in Lübeck (3-Tesla Siemens MAGNETOM Skyra magnetic resonance tomograph) using a 64-channel head coil. Two experimental tasks and the anatomical scan were conducted (see Supplement for details and scan acquisition parameters).

Functional MRI data analysis was performed using SPM12 (Wellcome Department of Imaging Neuroscience, London, UK; http://www.fil.ion.ucl.ac.uk/spm) implemented in Matlab R2019b (Mathworks Inc., Sherborn, MA, USA). Functional images were slice-timed and realigned to the first image to correct for head motion. Next, they were normalized to MNI space using the standard segmentation as implemented in SPM12 on the mean functional image and tissue probability maps. Normalization parameters were then applied to all images of the time series (resulting voxel size 3 × 3 × 3 mm). Finally, functional images were spatially smoothed with an 8 mm full width half maximum isotropic Gaussian kernel and high-pass filtered at 1/128 Hz to remove low-frequency drifts.

### Seed regions and functional connectivity analyses

Functional connectivity analyses were performed on a well-established reward processing paradigm (for details see Supplements). As in previous studies (32,33), a seed-based approach was used, from which all task-related activity was removed by regressing out hemodynamic responses induced by the task. Seed regions were the amygdala, ventral and dorsal striatum, thalamus, (dorsal and ventral) insula, ACC, and (anterior and posterior) DMN as alterations in AUD have been reported for these regions in the literature (see above). For details on the seed masks see Supplement.

To extract time series for the functional connectivity analysis from the seed regions, a first-level general linear model (GLM) was created for each participant, including predictor variables for the task (see Supplement for details), the six realignment parameters, and their temporal derivatives as nuisance regressors. Additionally, a scrubbing regressor was added to the GML, removing all volumes with a framewise displacement > 1 mm (see (34)). The time series of each ROI was then extracted using the first eigenvariate in SPM. Finally, new first-level GLMs were generated for every participant and every ROI, including the extracted time series of the respective ROI, all previously mentioned regressors, as well as regressors for white matter and cerebral spinal fluid signal. β-maps of each ROIs’ time series were analyzed at the second level (group-level) using a full factorial model in SPM12. To allow a comparison of group differences with the previous literature that did not perform a stress manipulation, a t-test comparing both groups only during the control task was also calculated. Whole-brain analyses were performed for each of the ROIs. An initial cluster-forming threshold of p < 0.001 (uncorrected) was used and statistical maps were then thresholded at a cluster-extent whole-brain false discovery rate (FDR)-corrected threshold of p < 0.05.

## Results

### Sample characteristics

Descriptive data on demographics, questionnaires, and baseline measures of pulse rate, cortisol, and negative affect can be found in Table 1.

**Table 1.** Sample characteristics.

|  | AUD<br>(N=34)<sup>a</sup> | Controls<br>(N=34)<sup>b</sup> | Group<br>comparisons |
| --- | --- | --- | --- |
| Gender | 97% m, 3% f | 94% m, 6% f |  |
| Age (years) | 42.41 ± 9.15 | 43.88 ± 9.31 |  |
| Duration of abstinence (days) | 14.76 ± 6.71 |  |  |
| Alcohol amount before withdrawal (g/day) | 301.88 ± 203.33 |  |  |
| BDI-FS | 3.98 ± 3.10 | 0.97 ± 1.31 | $F = 26.67, p < 0.001$ |
| TICS | 21.52 ± 8.41 | 10.33 ± 5.01 | $F = 44.44, p < 0.001$ |
| SRS | 56.75 ± 11.05 | 46.68 ± 7.23 | $F = 19.81, p < 0.001$ |
| Baseline pulse rate (bpm) | 79.80 ± 12.92 | 71.79 ± 12.45 | $t = 3.67, p < 0.001$ |
| Baseline cortisol (nmol/l) | 8.57 ± 5.34 | 8.32 ± 4.98 | $t = 0.28, p = 0.78$ |
| Baseline STAI-S | 23.36 ± 9.59 | 22.96 ± 8.47 | $t = 0.26, p = 0.79$ |

Values are given as mean and standard deviation if not specified otherwise. BDI-FS = Beck Depression Inventory- Fast Screening (56), the 7-item screening scale measures the severity of non-somatic depression symptomatology; TICS = Trier Inventory for Chronic Stress (57), values represent the 12-item screening scale as a global measure of experienced chronic stress; SRS = Stress Reactivity Scale (40), sum value as a measure of self-reported general stress reactivity; STAI-S = State part of the State-Trait-Anxiety Inventory (29).
<sup>a</sup>For one AUD subject, exact information about the duration of abstinence (between 10 and 40 days) and for two AUD subjects the exact amount of alcohol before withdrawal is missing, resulting in a sample size of N=33 and N=32 for these two variables, respectively.
<sup>b</sup>Pulse rate could not be recorded completely in case of two control subjects and one control subject did not fill in the BDI. Therefore, the sample size is reduced to N=32 and 33 for these two variables, respectively.

### Blunted physiological but increased affective stress responses in AUD

The three ANOVAs on changes in cortisol levels (AUCi) and difference scores of pulse rate (task phase - rest) and affect (STAI-S after - before the task) all revealed a significant main effect of *condition,* reflecting a greater increase in cortisol, pulse rate, and negative affect during/after the TSST compared to the control task, across all participants. In all three analyses, there was no main effect of *group*, but a significant interaction effect of *condition* and *group*: For cortisol and pulse rate, stress responses (in contrast to the control condition) were less pronounced in AUD compared to controls, but affective responses were stronger (Figure 1B).

### Physiological and affective stress responses correlate in healthy controls but not in AUD

Correlational analyses between stress-related changes in pulse rate, cortisol, and negative affect revealed significant positive associations between all three parameters in healthy controls, but no significant correlation in AUD (Figure 1C).

### Functional connectivity is decreased in AUD

Functional connectivity analyses revealed main effects of group for the connectivity of the dorsal striatum, ACC, and anterior and posterior DMN with several cortical and subcortical regions (Figure 2, Table 2), reflecting decreased connectivity in AUD. Additional t-tests comparing AUD and healthy controls on the control task only showed decreased connectivity of bilateral ventral and dorsal striatum, right dorsal insula, and anterior and posterior DMN with cortical and subcortical regions, specifically the striatum, cerebellum, and precuneus (Table S1).

**Figure 2.**
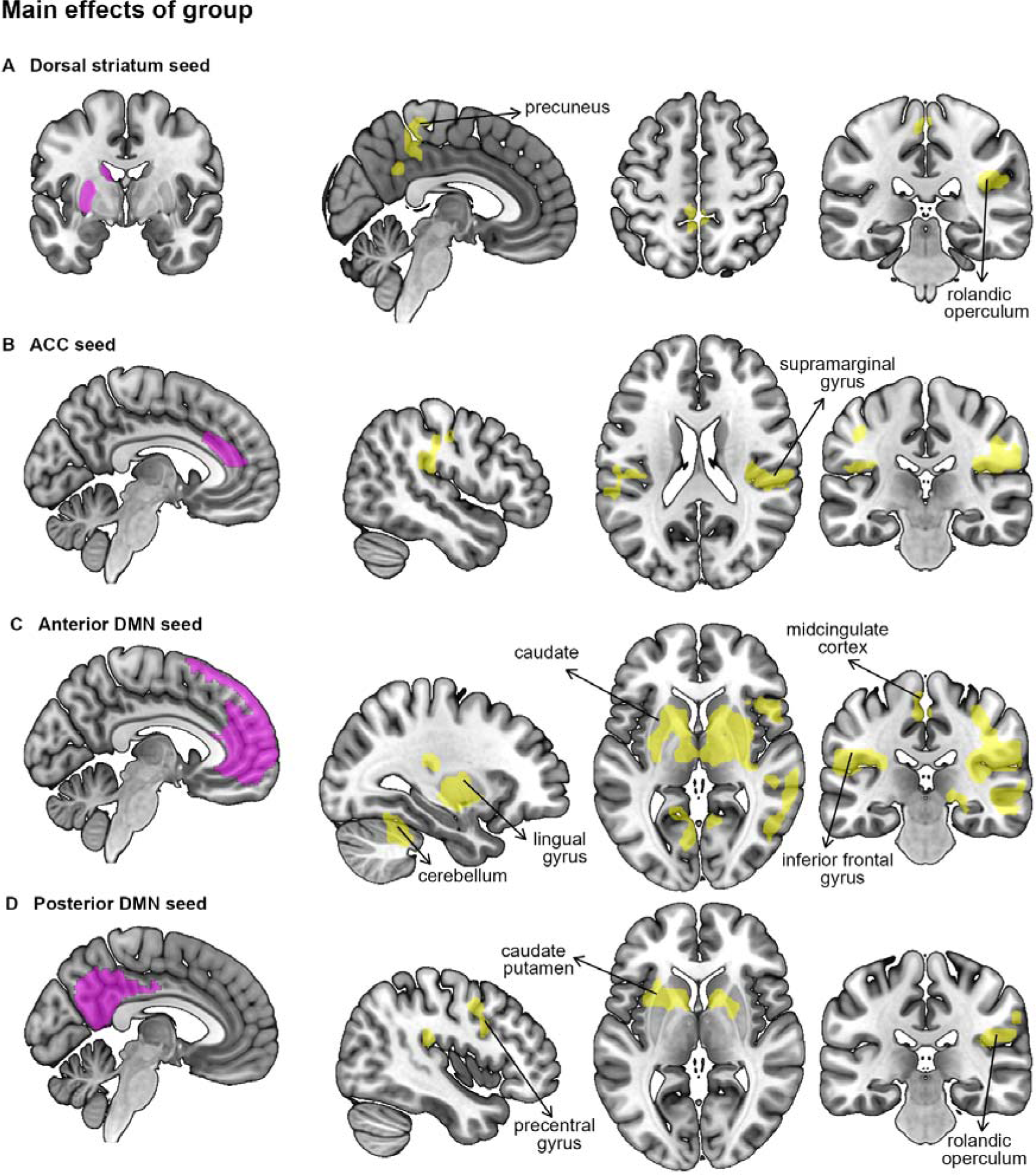
Main effects of group on functional connectivity of A) the dorsal striatum, B) the Anterior cingulate Cortex (ACC), C) the anterior Default Mode Network (DMN), and D) the posterior Mode Network. Functional connectivity of all depicted regions was significantly reduced in AUD compared to controls.

**Table 2.** Main effects of group on connectivity.

|  | side | MNI coordinates |  |  | Cluster size | F | p |
| --- | --- | --- | --- | --- | --- | --- | --- |
|  |  | x | y | z |  |  |  |
| **Seed: left dorsal striatum** |  |  |  |  |  |  |  |
| rolandic operculum, angular gyrus, sup. temp gyrus, supramarginal gyrus | R | 42 | -25 | 20 | 203 | 28.31 | 0.010 |
|  |  | 45 | -49 | 23 |  | 18.40 |  |
|  |  | 60 | -52 | 20 |  | 17.49 |  |
| precuneus, midcingulate cortex | L/R | 0 | -40 | 56 | 182 | 19.13 | 0.010 |
|  |  | -6 | -52 | 38 |  | 15.03 |  |
|  |  | 3 | -49 | 32 |  | 14.90 |  |
| **Seed: ACC** |  |  |  |  |  |  |  |
| rolandic operculum, supramarginal gyrus, postcentral gyrus, precentral gyrus | R | 42 | -25 | 20 | 294 | 26.88 | **0.001** |
|  |  | 57 | -25 | 23 |  | 18.16 |  |
|  |  | 63 | -19 | 35 |  | 17.28 |  |
| supramarginal gyrus, inferior parietal lobule | L | -57 | -40 | 32 | 227 | 18.13 | **0.002** |
|  |  | -45 | -52 | 53 |  | 17.17 |  |
|  |  | -51 | -46 | 47 |  | 17.14 |  |
| **Seed: anterior DMN** |  |  |  |  |  |  |  |
| caudate nucleus, rolandic operculum, supramarginal gyrus, putamen | L/R | 15 | 8 | 11 | 3238 | 37.33 | **<0.001** |
|  |  | 42 | -28 | 20 |  | 34.46 |  |
|  |  | -54 | -25 | 17 |  | 29.44 |  |
| fusiform gyrus, cerebellum, lingual gyrus | R | 30 | -40 | -22 | 508 | 27.84 | **<0.001** |
|  |  | 24 | -64 | -16 |  | 25.30 |  |
|  |  | 30 | -43 | -34 |  | 21.66 |  |
| cerebellum, hippocampus, lingual gyrus | L | -33 | -49 | -25 | 387 | 24.86 | **<0.001** |
|  |  | -18 | -37 | -1 |  | 21.82 |  |
|  |  | -6 | -76 | -1 |  | 19.62 |  |
|  |  | 57 | 26 | 14 |  | 13.12 |  |
| precuneus, midcingulate cortex,<br>paracentral lobule | L/R | -3 | -40 | 56 | 190 | 21.24 | 0.005 |
|  |  | 3 | -40 | 44 |  | 20.91 |  |
|  |  | -3 | -25 | 50 |  | 19.03 |  |
| inferior frontal gyrus | R | 54 | 20 | -1 | 131 | 27.95 | 0.021 |
|  |  | 57 | 11 | 8 |  | 22.64 |  |
| post. medial frontal, midcingulate<br>cortex | L/R | 3 | -10 | 65 | 103 | 17.95 | 0.044 |
|  |  | 0 | -7 | 47 |  | 17.65 |  |
| **Seed: posterior DMN** |  |  |  |  |  |  |  |
| caudate nucleus, putamen, insula<br>lobe | L | -9 | 8 | 11 | 233 | 23.60 | **0.002** |
|  |  | -24 | 11 | 2 |  | 20.56 |  |
|  |  | -33 | 5 | 11 |  | 15.65 |  |
| rolandic operculum, inferior<br>parietal lobule, supramarginal<br>gyrus | R | 42 | -28 | 20 | 138 | 20.70 | 0.016 |
|  |  | 57 | -37 | 47 |  | 17.50 |  |
|  |  | 63 | -22 | 29 |  | 14.78 |  |
| precentral gyrus, inferior frontal<br>gyrus | R | 45 | 2 | 44 | 126 | 18.64 | 0.016 |
|  |  | 51 | 5 | 35 |  | 17.12 |  |
|  |  | 48 | 11 | 26 |  | 16.20 |  |
| caudate nucleus, pallidum | R | 9 | 8 | 5 | 95 | 26.46 | 0.037 |
|  |  | 24 | 2 | 2 |  | 14.95 |  |

p < 0.001 (unc.), FDR, cluster-level, bold p-values mark results that survived Bonferroni correction to correct for multiple testing of the 15 seed regions (p < 0.05/15= 0.0033), ACC = Anterior Cingulate Cortex, DMN = Default Mode Network

### Interaction effects of group and stress on functional connectivity

Functional connectivity analyses did not reveal any main effect of stress in any of the ROIs. However, significant interaction effects of group and stress were found for functional connectivity of the left ventral and dorsal striatum with the right precuneus and superior parietal lobule, of the left dorsal striatum with the right cerebellum and midcingulate cortex, of the right ventral insula with the cerebellum, and the posterior DMN with the left putamen and thalamus (Table 3, Figure 3).

**Figure 3.**
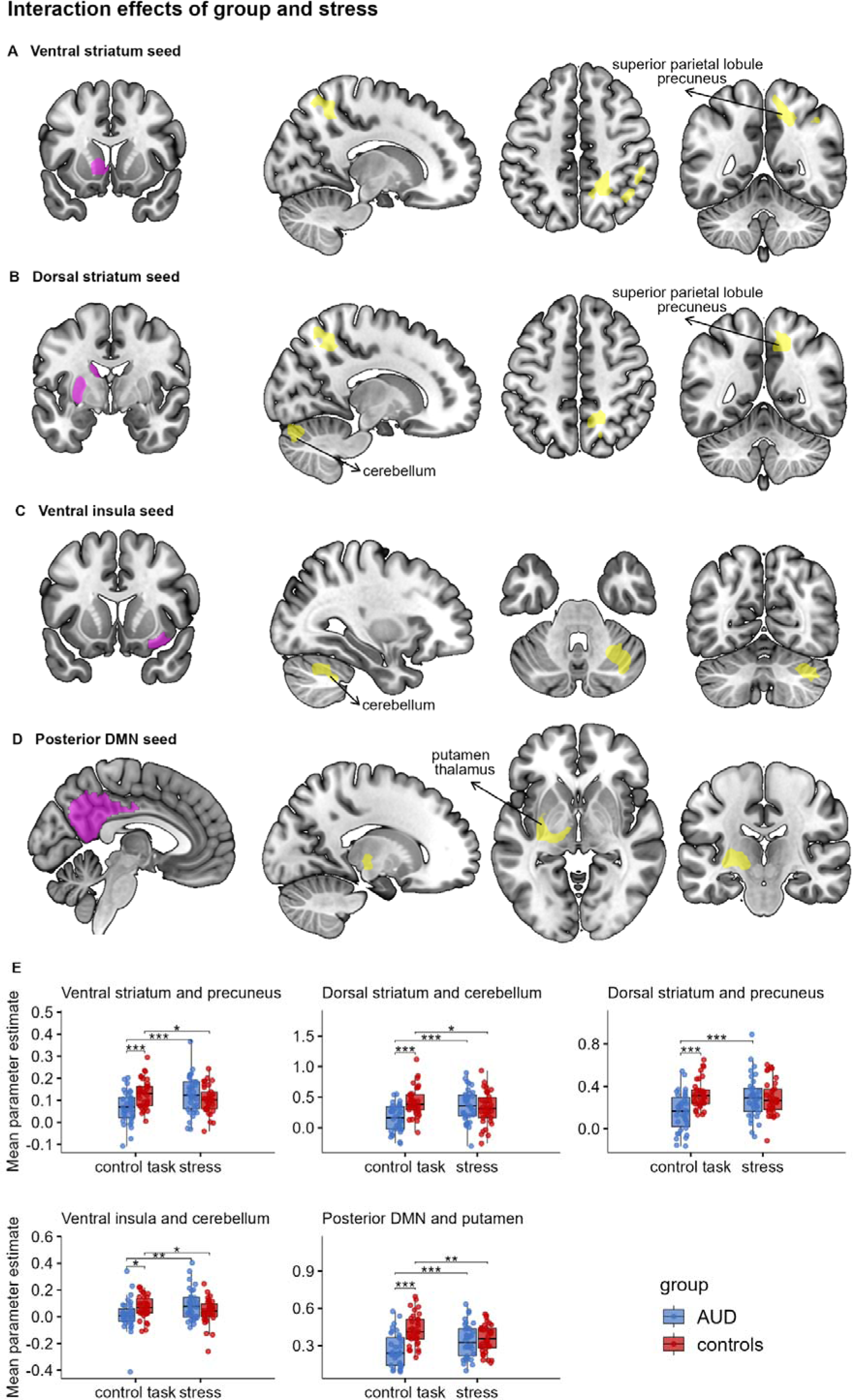
Group x stress interaction effects on functional connectivity of A) the ventral striatum seed, B) the dorsal striatum seed, C) the ventral insula DMN seed, and D) the posterior DMN seed. E) Mean parameter estimates for patients with AUD and healthy controls of the clusters with significant group x stress interactions show that functional connectivity is decreased in AUD on the control day. After stress, functional connectivity increases in AUD and decreases in controls leading to comparable results.

**Table 3.**
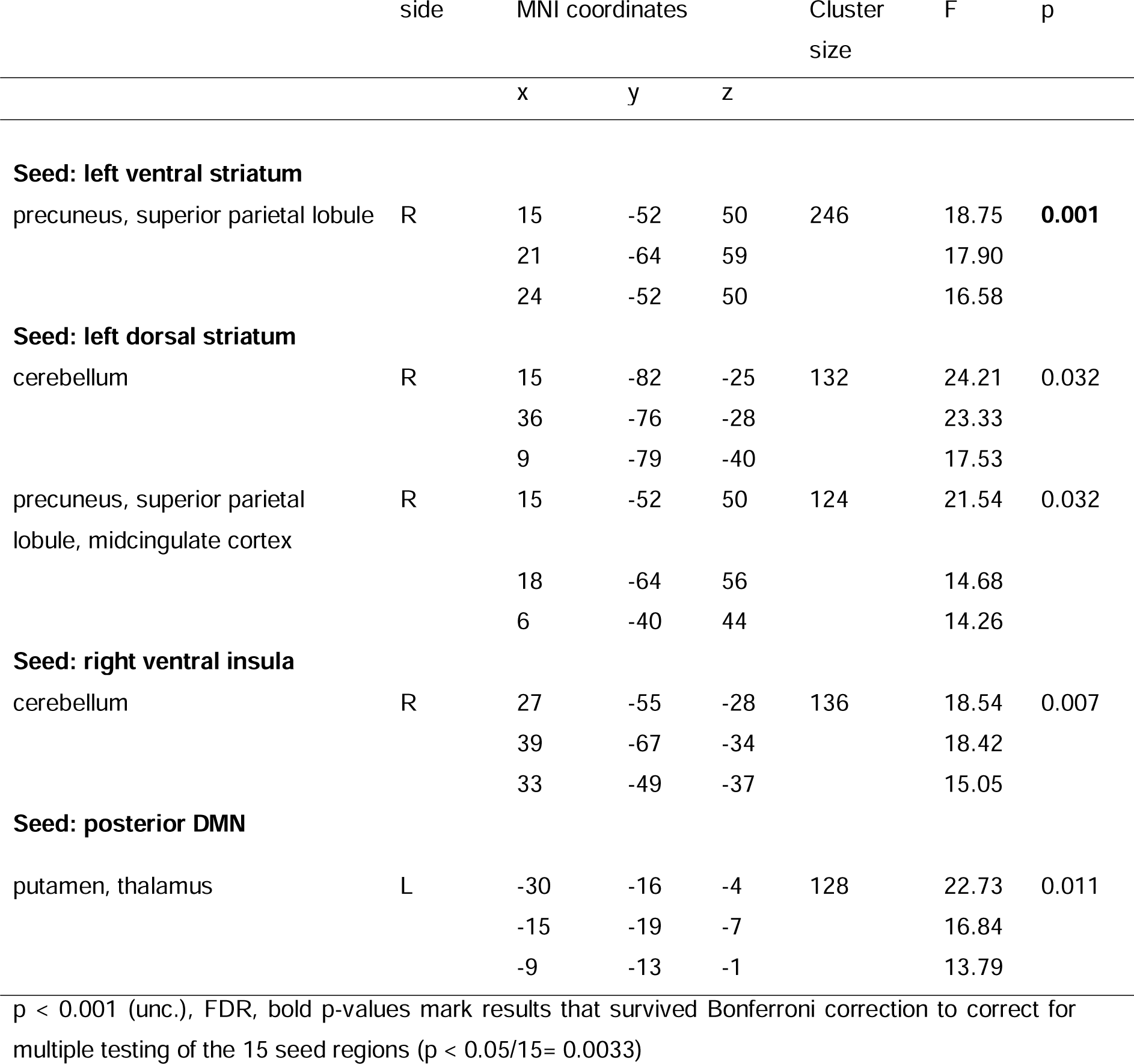
Interaction effects of group and stress on connectivity.

## Discussion

The present study investigated physiological and affective responses to acute stress along with functional brain connectivity in AUD. Altered stress responses were found for all stress parameters (cortisol, pulse rate, and negative affect) in AUD compared to controls. In parallel, stress effects on functional connectivity also differed between groups.

Importantly, the TSST induced increases in cortisol, pulse rate, and negative affect across groups suggesting successful stress induction. Furthermore, group comparisons of baseline levels of cortisol and pulse rate replicated previous findings in the literature: We found no group difference in basal cortisol which is consistent with reports that although cortisol levels are elevated during intoxication and acute withdrawal, they decline during abstinence and are no longer different from those of healthy subjects in most studies (see (6) for review). Baseline pulse rate was elevated in AUD in the present study, consistent with previous findings of elevated basal heart or pulse rate in the first weeks of abstinence (7–10). These alterations are assumed to be a consequence of chronic alcohol consumption (7–10): Acute alcohol intake affects the cardiovascular system via centrally mediated sympathetic activity and abnormalities of cardiovascular regulation appear to persist into recovery (35).

Responses to acute stress were altered in AUD for all measured parameters, but the effects differed in their direction. Phasic cortisol responses after stress induction were significantly lower in AUD confirming our hypothesis and replicating previous findings of blunted cortisol responses to stress in AUD (see (12) for review). The design of the present study does not allow conclusions as to whether the alterations found are a consequence of alcohol consumption or precede it. However, previous studies indicate that blunted cortisol responses are due to a dysregulation of glucocorticoid signaling following repetitive activation of the HPA axis by chronic alcohol intake (36) and that they normalize with long-term abstinence (12). Animal studies suggest that corticotropin-releasing factor (CRF) may play a critical role in this process (37,38). For pulse rate, we also found smaller differences between the stress and control tasks in AUD as compared to controls. However, this interaction effect was not due to a blunted response during stress, but rather to an increased response in AUD during the control condition. This finding may explain why many previous studies - at least those using social stress tasks - have found no group differences (see (6) for review). These studies usually did not include a non-stressful control task and only compared pulse changes in response to stress between groups (except (39) who also found an interaction effect). This comparison would not have shown a group difference in our study either. These results suggest that it may be important to include a control condition in future studies to better understand stress-related pulse or heart rate changes in AUD.

In contrast to the findings for cortisol and pulse rate, stress-related negative affect was increased in AUD compared to controls replicating previous studies (7,17,18). Consistent with this, SRS scores, which represent affective stress reactivity as a trait (40), were also significantly elevated in AUD. Taken together, these findings suggest a dissociation between subjectively experienced distress (increased) and the physiological stress response (decreased HPA and sympathetic responses compared to the control condition) in AUD. This dissociation is further corroborated by the finding that stress-related changes in pulse rate, cortisol, and negative affect were correlated in healthy subjects but not in AUD. We assume that a dysregulation of the physiological stress systems in AUD underlies this finding. Interestingly, postmortem analyses of gene expression in the hippocampus have shown that pathways involved in stress responses are mostly increased in AUD (41), which may indicate that blunted physiological stress responses do not reflect decreased activity of brain stress systems (37). Furthermore, the increased affective stress response may also be due to a dysregulation of brain stress systems: In rats, chronic alcohol consumption has been shown to lead to heightened activity of the CRF system in the central amygdala in response to stress (42) and alterations in the CRF system have been suggested to underlie enhanced behavioral stress responses (43). These alterations can persist after longer periods of abstinence (37). While these findings suggest that heightened affective stress responses are a result of chronic alcohol consumption, there may also be trait factors such as negative emotionality, that precede the onset of substance abuse (44). Longitudinal studies are needed to clarify whether the increased affective stress responses are a consequence of alcohol consumption or a trait and risk factor.

Neuroimaging analyses revealed decreased functional connectivity of the left dorsal striatum, ACC, and anterior and posterior DMN with several cortical and subcortical regions in AUD across both test days. A separate analysis of only the control day additionally revealed reduced connectivity of the bilateral ventral striatum, right dorsal striatum, and right dorsal insula with various regions (mainly striatum, cerebellum, precuneus). The largest alterations of connectivity were found for the seeds of the anterior and posterior DMN, supporting the notion that this network plays an important role in substance use disorders (SUD) (27,45). Several of our findings replicate previous results of resting-state fMRI research in AUD, including decreases in connectivity between the anterior and posterior DMN (27), between the anterior DMN and the putamen (25), between the ventral striatum and angular gyrus (26), between the caudate and superior frontal gyrus as well as posterior DMN (26), and between the insula and cerebellum (46). However, there were also findings in the literature that contradict our results such as increased connectivity between the striatum and superior frontal gyrus, inferior parietal gyrus, or cerebellum (22,23). A possible explanation for this could be related to the analytical approach of the present study. We assessed functional connectivity during the processing of a reward task. Although the effects of the task were adjusted for (see (29)), it may have affected the connectivity of the striatum, a central structure of the reward system. There was, however, also a great heterogeneity in the results of previous resting-state fMRI research, possibly related to variations in the duration of abstinence and other sample variables.

Previous research has reported altered connectivity of the amygdala, thalamus, and DMN (PCC) after stress induction in the healthy population (21). In the present study comprising healthy individuals and patients with AUD, we did not observe any significant effect of stress in any of these ROIs across all subjects. However, interaction effects of stress and group were found for the left ventral and dorsal striatum, right ventral insula, and posterior DMN seed, reflecting decreased connectivity on the control day but normal connectivity after stress in AUD.

This effect was found for the connectivity of the left ventral and dorsal striatum seed with the right superior parietal lobule and precuneus and similarly for the posterior DMN seed (encompassing the precuneus) with the putamen. It has been previously reported that the striatum is functionally connected to the precuneus/posterior DMN and superior parietal regions in healthy subjects (47) and that resting-state connectivity between the posterior DMN and striatum is decreased in drug-addicted individuals (27). Our results now suggest that in AUD, this connection is more strongly involved after acute stress (increase in connectivity after stress while controls show a decrease). It is unclear what mechanisms underlie this effect. It has been suggested that altered connectivity between the DMN and subcortical areas enhances negative emotions in SUD, but the role of specific connectivity tracts has not been investigated (27). Alterations in the dopamine system in AUD may play a role here. Chronic drug administration alters striatal dopamine signaling and studies in healthy subjects and animals suggest that changes in dopamine modulation might affect (de)activation of the DMN (27). The dopaminergic system is also involved in stress: Acute stress induces CRF release which leads to dopamine release in the striatum (48). However, if the stress axes are dysregulated due to chronic substance consumption, the ability of CRF to modulate dopamine levels may be abolished, as has already been postulated for chronic stress (48). These changes in dopamine modulation may be a potential mechanism by which alterations in connectivity between the striatum and posterior DMN after acute stress may occur in AUD. They may also underlie the interaction effect found for the connectivity between the posterior DMN and the thalamus, as nuclei within the thalamus are innervated by dopamine and CRF among other neurotransmitters, and are closely connected to the striatum (49–51), which has been proposed to be linked to stress-associated changes in dopamine tone (52).

Interaction effects of stress and group in the same direction were also found for the connectivity of both the dorsal striatum seed and the ventral insula seed with the cerebellum. A recent study also reported decreased connectivity between the insula and the cerebellum in AUD (53). In addition, several studies have implicated the cerebellum in the stress response (see (54) for review). It has been suggested that the cerebellum may play an important role in stress-associated neurobehavioral effects because it expresses the cellular machinery necessary to process neurochemical mediators such as glucocorticoids and CRF among others (54). Thus, acute stress may have an altered effect on cerebellar connectivity in AUD via these systems.

Two limitations should be considered when interpreting the study results. First, as part of a broader project, blood samples were collected which already caused stress in some individuals, as evidenced by an increase in cortisol levels from T0 to T1 in a sub-sample. Although there was still a rest period between the blood draw and the time when the pre-stress data collection, this may have influenced the results for cortisol in particular, as there is a time lag between the rise and fall of salivary cortisol levels. Second, cortisol from saliva samples does not necessarily reflect brain glucocorticoid concentrations. Research in rodents has shown that withdrawal from long-term alcohol consumption was related to increases in glucocorticoid concentrations in specific brain regions, while there was no change in plasma levels (55). Thus, the relationship between systemic and brain glucocorticoids appears to be complex, suggesting caution when interpreting the results.

In conclusion, our findings provide evidence for a dissociation between physiological and affective stress responses in AUD, as well as stress-related alterations in functional connectivity of striatal, insular, cerebellar, and default mode regions. These results suggest a complex interplay between chronic alcohol use and acute stress regulation at physiological, affective, and neural levels, which may be important to consider for therapeutic approaches.

## Acknowledgements

This work was supported by the Else Kröner-Fresenius-Stiftung (to LR, Grant No. 2018_A26). YS and JV were funded by the Studienstiftung des Deutschen Volkes. YS has previously presented data from this study at the 2. Deutscher Psychotherapie Kongress, Berlin, Germany (May 2023) and at the Psychologie und Gehirn conference, Tübingen, Germany (June 2023).

## Footnotes

1 Please note that also blood samples were taken throughout the test day which are not relevant for this study. See Supplements for details.

## References

1. Milivojevic V, Sinha R (2018): Central and Peripheral Biomarkers of Stress Response for Addiction Risk and Relapse Vulnerability. Trends Mol Med 24: 173–186.

2. Adinoff B, Ruether K, Krebaum S, Iranmanesh A, Williams MJ (2003): Increased salivary cortisol concentrations during chronic alcohol intoxication in a naturalistic clinical sample of men. Alcohol Clin Exp Res 27: 1420–1427.

3. Kutscher S, Heise DJ, Banger M, Saller B, Michel MC, Gastpar M, et al. (2002): Concomitant endocrine and immune alterations during alcohol intoxication and acute withdrawal in alcohol-dependent subjects. Neuropsychobiology 45: 144–149.

4. Price JL, Nixon SJ (2021): Retrospective Hair Cortisol Concentrations from Pretreatment to Early Recovery in Alcohol Use Disorder. Alcohol Alcohol 56: 181–184.

5. Stephens MAC, Wand G (2012): Stress and the HPA axis: role of glucocorticoids in alcohol dependence. Alcohol Res 34: 468–483.

6. Chen K, Hollunder B, Garbusow M, Sebold M, Heinz A (2020): The physiological responses to acute stress in alcohol-dependent patients: A systematic review. Eur Neuropsychopharmacol 41: 1–15.

7. Sinha R, Fox HC, Hong KA, Bergquist K, Bhagwagar Z, Siedlarz KM (2009): Enhanced negative emotion and alcohol craving, and altered physiological responses following stress and cue exposure in alcohol dependent individuals. Neuropsychopharmacology 34: 1198–1208.

8. Tolic I, Soyka M (2018): Stressreagibilität bei Alkoholabhängigen unter Berücksichtigung von Abstinenzdauer und Krankheitsschwere. Fortschr Neurol Psychiatr 86: 356–367.

9. Demmel R, Rist F, Olbrich R (2000): Autonomic reactivity to mental stressors after single administration of lorazepam in male alcoholics and healthy controls. Alcohol Alcohol 35: 617–624.

10. Seo D, Lacadie CM, Tuit K, Hong K-I, Constable RT, Sinha R (2013): Disrupted ventromedial prefrontal function, alcohol craving, and subsequent relapse risk. JAMA Psychiatry 70: 727–739.

11. Ralevski E, Petrakis I, Altemus M (2019): Heart rate variability in alcohol use: A review. Pharmacol Biochem Behav 176: 83–92.

12. Dunne N, Ivers J-H (2023): HPA axis function in alcohol use disorder: A systematic review and meta-analysis. Addiction Neuroscience 8: 100114.

13. Junghanns K, Tietz U, Dibbelt L, Kuether M, Jurth R, Ehrenthal D, et al. (2005): Attenuated salivary cortisol secretion under cue exposure is associated with early relapse. Alcohol Alcohol 40: 80–85.

14. Adinoff B, Junghanns K, Kiefer F, Krishnan-Sarin S (2005): Suppression of the HPA axis stress-response: implications for relapse. Alcohol Clin Exp Res 29: 1351–1355.

15. Junghanns K, Backhaus J, Tietz U, Lange W, Bernzen J, Wetterling T, et al. (2003): Impaired serum cortisol stress response is a predictor of early relapse. Alcohol Alcohol 38: 189–193.

16. Starcke K, van Holst RJ, van den Brink W, Veltman DJ, Goudriaan AE (2013): Physiological and endocrine reactions to psychosocial stress in alcohol use disorders: duration of abstinence matters. Alcohol Clin Exp Res 37: 1343–1350.

17. Romero-Martínez Á, Vitoria-Estruch S, Moya-Albiol L (2019): Emotional and autonomic dysregulation in abstinent alcoholic men: An idiosyncratic profile? Alcohol 77: 155–162.

18. Bernardy NC, King AC, Lovallo WR (2003): Cardiovascular responses to physical and psychological stress in female alcoholics with transitory hypertension after early abstinence. Alcohol Clin Exp Res 27: 1489–1498.

19. van den Heuvel MP, Hulshoff Pol HE (2010): Exploring the brain network: a review on resting-state fMRI functional connectivity. Eur Neuropsychopharmacol 20: 519–534.

20. Wade NE, Padula CB, Anthenelli RM, Nelson E, Eliassen J, Lisdahl KM (2017): Blunted amygdala functional connectivity during a stress task in alcohol dependent individuals: A pilot study. Neurobiol Stress 7: 74–79.

21. van Oort J, Tendolkar I, Hermans EJ, Mulders PC, Beckmann CF, Schene AH, et al. (2017): How the brain connects in response to acute stress: A review at the human brain systems level. Neurosci Biobehav Rev 83: 281–297.

22. Kohno M, Dennis LE, McCready H, Hoffman WF (2017): Executive Control and Striatal Resting-State Network Interact with Risk Factors to Influence Treatment Outcomes in Alcohol-Use Disorder. Front Psychiatry 8: 182.

23. Camchong J, Stenger A, Fein G (2013): Resting-state synchrony in long-term abstinent alcoholics. Alcohol Clin Exp Res 37: 75–85.

24. Liu J, Cai W, Zhao M, Cai W, Sui F, Hou W, et al. (2019): Reduced resting-state functional connectivity and sleep impairment in abstinent male alcohol-dependent patients. Hum Brain Mapp 40: 4941–4951.

25. Wang J, Fan Y, Dong Y, Ma M, Ma Y, Dong Y, et al. (2016): Alterations in Brain Structure and Functional Connectivity in Alcohol Dependent Patients and Possible Association with Impulsivity. PLoS One 11: e0161956.

26. Müller-Oehring EM, Jung Y-C, Pfefferbaum A, Sullivan EV, Schulte T (2015): The Resting Brain of Alcoholics. Cereb Cortex 25: 4155–4168.

27. Zhang R, Volkow ND (2019): Brain default-mode network dysfunction in addiction. Neuroimage 200: 313–331.

28. Kirschbaum C, Pirke KM, Hellhammer DH (1993): The “Trier Social Stress Test”--a tool for investigating psychobiological stress responses in a laboratory setting. Neuropsychobiology 28: 76–81.

29. Grimm J (2009): State-Trait-Anxiety Inventory Nach Spielberger. Deutsche Lang-Und Kurzversion. Methodenforum Der Universität Wien. MF-Working Paper 2009/02.

30. Pruessner JC, Kirschbaum C, Meinlschmid G, Hellhammer DH (2003): Two formulas for computation of the area under the curve represent measures of total hormone concentration versus time-dependent change. Psychoneuroendocrinology 28: 916–931.

31. Diedenhofen B, Musch J (2015): cocor: A Comprehensive Solution for the Statistical Comparison of Correlations. PLoS ONE, vol. 10. Public Library of Science, p e0121945.

32. Esslinger C, Walter H, Kirsch P, Erk S, Schnell K, Arnold C, et al. (2009): Neural Mechanisms of a Genome-Wide Supported Psychosis Variant. Science 324: 605.

33. Paulus FM, Krach S, Bedenbender J, Pyka M, Sommer J, Krug A, et al. (2013): Partial support for ZNF804A genotype-dependent alterations in prefrontal connectivity. Hum Brain Mapp 34: 304–313.

34. Power JD, Mitra A, Laumann TO, Snyder AZ, Schlaggar BL, Petersen SE (2014): Methods to detect, characterize, and remove motion artifact in resting state fMRI. Neuroimage 84: 320–341.

35. Irwin MR, Ziegler M (2005): Sleep deprivation potentiates activation of cardiovascular and catecholamine responses in abstinent alcoholics. Hypertension 45: 252–257.

36. Edwards S, Little HJ, Richardson HN, Vendruscolo LF (2015): Divergent regulation of distinct glucocorticoid systems in alcohol dependence. Alcohol 49: 811–816.

37. Fosnocht AQ, Briand LA (2016): Substance use modulates stress reactivity: Behavioral and physiological outcomes. Physiol Behav 166: 32–42.

38. Zorrilla EP, Valdez GR, Weiss F (2001): Changes in levels of regional CRF-like-immunoreactivity and plasma corticosterone during protracted drug withdrawal in dependent rats. Psychopharmacology 158: 374–381.

39. Panknin TL, Dickensheets SL, Nixon SJ, Lovallo WR (2002): Attenuated heart rate responses to public speaking in individuals with alcohol dependence. Alcohol Clin Exp Res 26: 841–847.

40. Schulz P, Jansen LJ, Schlotz W (2005): Stressreaktivität: Theoretisches Konzept und Messung. Diagnostica 51: 124–133.

41. McClintick JN, Xuei X, Tischfield JA, Goate A, Foroud T, Wetherill L, et al. (2013): Stress-response pathways are altered in the hippocampus of chronic alcoholics. Alcohol 47: 505–515.

42. Retson TA, Reyes BA, Van Bockstaele EJ (2015): Chronic alcohol exposure differentially affects activation of female locus coeruleus neurons and the subcellular distribution of corticotropin releasing factor receptors. Prog Neuropsychopharmacol Biol Psychiatry 56: 66–74.

43. Valdez GR, Zorrilla EP, Roberts AJ, Koob GF (2003): Antagonism of corticotropin-releasing factor attenuates the enhanced responsiveness to stress observed during protracted ethanol abstinence. Alcohol 29: 55–60.

44. Measelle JR, Stice E, Springer DW (2006): A prospective test of the negative affect model of substance abuse: moderating effects of social support. Psychol Addict Behav 20: 225–233.

45. Taebi A, Becker B, Klugah-Brown B, Roecher E, Biswal B, Zweerings J, Mathiak K (2022): Shared network-level functional alterations across substance use disorders: A multi-level kernel density meta-analysis of resting-state functional connectivity studies. Addict Biol 27: e13200.

46. Halcomb ME, Chumin EJ, Goñi J, Dzemidzic M, Yoder KK (2019): Aberrations of anterior insular cortex functional connectivity in nontreatment-seeking alcoholics. Psychiatry Res Neuroimaging 284: 21–28.

47. Di Martino A, Scheres A, Margulies DS, Kelly AMC, Uddin LQ, Shehzad Z, et al. (2008): Functional connectivity of human striatum: a resting state FMRI study. Cereb Cortex 18: 2735–2747.

48. Kumar P, Berghorst LH, Nickerson LD, Dutra SJ, Goer FK, Greve DN, Pizzagalli DA (2014): Differential effects of acute stress on anticipatory and consummatory phases of reward processing. Neuroscience 266: 1–12.

49. García-Cabezas MA, Rico B, Sánchez-González MA, Cavada C (2007): Distribution of the dopamine innervation in the macaque and human thalamus. Neuroimage 34: 965– 984.

50. Hsu DT, Kirouac GJ, Zubieta J-K, Bhatnagar S (2014): Contributions of the paraventricular thalamic nucleus in the regulation of stress, motivation, and mood. Front Behav Neurosci 8: 73.

51. Huang AS, Mitchell JA, Haber SN, Alia-Klein N, Goldstein RZ (2018): The thalamus in drug addiction: from rodents to humans. Philos Trans R Soc Lond B Biol Sci 373. 10.1098/rstb.2017.0028

52. Parsons MP, Li S, Kirouac GJ (2007): Functional and anatomical connection between the paraventricular nucleus of the thalamus and dopamine fibers of the nucleus accumbens. J Comp Neurol 500: 1050–1063.

53. Manuweera T, Kisner MA, Almira E, Momenan R (2022): Alcohol use disorder-associated structural and functional characteristics of the insula. J Neurosci Res 100: 2077–2089.

54. Moreno-Rius J (2019): The cerebellum under stress. Front Neuroendocrinol 54: 100774.

55. Little HJ, Croft AP, O’Callaghan MJ, Brooks SP, Wang G, Shaw SG (2008): Selective increases in regional brain glucocorticoid: a novel effect of chronic alcohol. Neuroscience 156: 1017–1027.

56. Beck AT, Steer RA, Brown GK (2013): Beck Depressions-Inventar-FS. Deutsche Bearbeitung von Sören Kliem & Elmar.

57. Schulz P, Schlotz W, Becker P, Schulz P, Schlotz W, Becker P (2004): Trierer Inventar zum Chronischen Stress (TICS) [Trier Inventory for Chronic Stress (TICS)]. Retrieved February 15, 2023, from https://eprints.soton.ac.uk/50017/

