## Supplemental Material for "Altered physiological, affective, and functional connectivity responses to acute stress in patients with alcohol use disorder"

### **Supplemental information**

### **Sample: Recruitment and inclusion/ exclusion criteria**

Patients were recruited through the Department of Psychiatry and Psychotherapy in Lübeck and the AMEOS clinic in Lübeck, Germany. Healthy controls were recruited via flyers, letters, and online advertisements.

Participants were required to meet the following criteria (which were in part due to other research questions in the project concerning the investigation of inflammatory parameters): age between 18 and 60 years, sufficient knowledge of the German language, scanner compatibility, no neurological disorders or brain injuries, no medication that can have an effect on the HPA-axis or dopamine system in the past two weeks, no acute infections, no vaccination in the past 3 weeks, and no chronic inflammatory diseases. Patients additionally had to meet the following criteria: diagnosis of alcohol use disorder according to the Diagnostic and Statistical Manual of Mental Disorders fifth edition (DSM-V), alcohol abstinence for at least 10 days, no comorbid diagnosis of major depression, antisocial or borderline personality disorders, social anxiety, psychosis, or acute suicidality. Further inclusion criteria for control participants were: no current or past psychiatric disorders, no alcohol misuse classified as ≥ 8 points in the alcohol use disorder identification test (1), no consumption of drugs (except for alcohol, nicotine, and irregular use of cannabis) in the past year and no history of regular intake/dependency of any drug except nicotine.

### **Sample: Dropouts and exclusions**

Two patients and one control subject dropped out of the study. In addition, four patients and two control subjects had to be excluded from the analyses because of technical problems or excessive head movement during MRI scanning on one of the two testing days. Furthermore, two patients and one control subject did not participate in MRI scanning for safety reasons and took only part in the other measurements. Please note that the current study is part of a broader project and data of these subjects can be used to examine other research questions of the project. In this study, however, only subjects with usable MRI data sets of both testing days were included.

### **Procedure: Interval between the two test days**

The two days of testing took place at a maximum of 10 days apart for all but one control subject. In the case of this individual, measurements could not be performed on the originally scheduled second test day because the participant was not well. Due to his working schedule, he was only able to return for the second test day after 48 days. Excluding this participant from the analyses had no substantial impact on the results. The only two changes in the statistics were that in the ANOVA on negative affect the main effect of group now reached significance (*F*(1, 65) = 4.10, *p* = .047, *η_p_^2^*= .06) - what had previously been slightly above the threshold - and that in the functional connectivity analysis an interaction effect between the left ventral striatum and the cerebellum now also became significant.

### **Procedure: Blood samples**

Blood samples were also collected as part of a larger project, but these are not analyzed in this study. Before the SARS-CoV-19 pandemic, blood samples were taken at the beginning and at three time points throughout the test day (after the TSST/control task, directly after MRI scanning and at the end of the test day). After the pause (see below), they were only taken at the beginning and end of the test day.

### **Procedure: Data collection before and during the SARS-CoV-19 pandemic**

Due to the SARS-CoV-19 pandemic, data collection was temporarily paused between March 2020 and October 2021, resulting in a subset of participants being recruited before (N=49) and others during the pandemic at a time when the risk of infection for study personnel and participants was more manageable because of vaccinations and testing (N=31). After the break, there were some procedural changes, for example a SARS-CoV-19 rapid test was conducted upon arrival of the participant and face masks were worn during the test day. In addition, for organizational reasons, it was decided to draw blood at only two time points instead of four, using venipuncture instead of an indwelling venous catheter. As it cannot be ruled out that the changes in the study protocol after the break, as well as general societal changes due to the pandemic and associated stress, could affect the study data, we reran all analyses with a covariate distinguishing between measurements made before and during the pandemic. This yielded no substantial influence on the results. There was only one minor change in the statistics after inclusion of the covariate: the main effect of group in the ANOVA on negative affect reached significance (*F*(1, 65) = 5.05, *p* = .028, *η_p_^2^*= .07), which had previously been slightly above the threshold.

### **Stress induction: Trier Social Stress Test and control session**

To induce acute psychosocial stress in participants, the Trier Social Stress Test (2) was used - a reliable and widely used tool for investigating acute psychosocial stress under experimental conditions (3). The TSST is a standardized protocol that includes a 10-minute anticipation phase, followed by a 10-minute test phase. According to the protocol, participants are first introduced to the tasks they are required to complete, which includes delivering a free speech of five-minute duration and performing a mental arithmetic task in front of a jury. The jury consists of three persons, who are briefly introduced to the participant in a second room to increase psychosocial stress. For the anticipation phase, participants return to the first room and are provided with pen and paper to prepare for the upcoming task while the experimenter waits outside. After ten minutes, the experimenter takes the participant back to the jury room and the test phase starts, consisting of the five-minute speech, in which they should convince the jury that they are the perfect candidate for a fictional vacant position and five minutes of mental arithmetic, in which the participant serially has to subtract 13 from the number 1022. In the current study, the protocol of the TSST was followed, with only minor deviations: the experimenter did not leave the room during the test phase but was part of the jury and no microphone was used. As a cover story, participants were informed that the task was designed to test their verbal and mathematical abilities.

In addition to the stress induction procedure, a control session (similar as in (4)) was included on a separate day, in which participants only read out written texts (instead of a free speech as in (4)) and performed easy mental arithmetic in the absence of a jury. This was very similar to the stress induction procedure (both took place in the same rooms, participants were in standing position during the task phase, and both involved verbal and mathematical tasks) but should not elicit psychosocial stress associated with performance evaluation. Here, participants were informed that the purpose of the tasks was to assess the physiological effects of mental effort by measuring their pulse rate as part of the cover story (when the task was performed on the first test day).

### **Cortisol: time points of saliva collection and laboratory analysis**

Saliva samples were collected at five timepoints during the test day: approximately 10 minutes after the arrival of the participant (T0), immediately before the introduction of the stress/control task (T1), after the stress/control task, which was on average 13 minutes after the onset of the task phase (T2), immediately before the MRI session, which was on average 29 minutes after the onset of the task phase of the stress/control task (T3), and after the MRI session, which was on average 78 minutes after the onset of the task phase of the stress/control task (T4).

After collection, samples were frozen and stored at -20 degrees Celsius until they were sent to the lab of Dresden LabService GmbH. After thawing, salivettes were centrifuged at 3000 rpm for five minutes, which resulted in a clear supernatant of low viscosity. Salivary concentrations were measured by commercially available chemiluminescence immunoassay with high sensitivity (IBL International, Hamburg, Germany). The intra- and inter-assay coefficients of variance were below 9%.

**Pulse rate**

Mean pulse rate scores for three phases of recording were computed: 1) the resting phase (baseline, the last 10 minutes were used for the analysis), 2) the anticipation phase of the TSST/control task (10 minutes), and 3) the actual task phase (10 minutes).

**Negative affect ratings**

To assess affective changes after the stress induction, the state part of the State-Trait-Anxiety Inventory (STAI-S) short version (5) was administered twice on both test days: once right before the introduction of the TSST or control task and once immediately after the task phase. The STAI-S short version comprises ten self-report items on an eight-point Likert scale which measure temporary subjective feelings of nervousness, tension, and worry. It refers to how a person is feeling in that moment and thus is often used in experimental stress research. Three subjects had a missing value which was replaced by the mean value of the other nine items for the analyses.

**Other questionnaires**

Furthermore, participants filled in questionnaires that measure more persistent variables and were therefore collected only once. On the first test day, these included the Beck Depression Inventory (BDI-II) (6) to assess depressive symptoms in the last week. Since alcohol withdrawal can be accompanied by numerous somatic complaints, we decided to consider only non-somatic criteria for assessing depressive symptoms as recommended for patients with medical problems including addictive disorders (BDI-FS; (7)). On the second day, participants completed the Stress Reactivity Scale (SRS; (8)), which measures stress reactivity defined as a trait that relatively stable constitutes the individual affective reaction to stressors as well as the screening scale of the Trier Inventory for Chronic Stress (TICS; (9)), which assesses experienced stress in the last three months. Thus, all questionnaires that made reference to stress were given only at the end of the second test day to ensure that participants could not already identify stress as a subject of the study prior to stress induction. For the analyses, the sum value of the SRS as a measure of general stress reactivity and the 12-item screening scale of the TICS as a global measure of experienced chronic stress were calculated according to the manuals. Missing values were replaced by the mean.

### **fMRI: Acquisition parameters**

A single-shot gradient-recalled echo-planar imaging (GRE-EPI) sequence was used to acquire functional images (repeat time (TR) = 1330 ms, echo time (TE) = 25 ms, flip angle = 70°, field of view (FoV) = 560x560). Additionally, an anatomical T1 image was obtained for each participant, which was acquired using an MP-RAGE sequence (TR = 1900 ms, TE = 2.44 ms, TI = 900 ms, flip angle = 9°, resolution 1×1×1 mm3, FOV = 192×256×256 mm3; acquisition time = 4.5 minutes).

### **fMRI: Description of the fMRI tasks**

During the MRI scan, participants performed two tasks and an anatomical scan. The first task was an adaptation of the well-established monetary incentive delay/ social incentive delay (MID/SID) paradigm (10–12), which is used to examine participants’ neural reactions to anticipated monetary and social rewards. In this task, visual cues indicate whether a reward can be won. Subsequently, a target stimulus is presented, to which the participant responds as quickly as possible by pressing a button. If the reaction was fast enough (adapted to individual response times), a reward is displayed afterwards with images of smiling faces used as social rewards and images of small amounts of money used in the monetary condition. If the reaction was too slow, only blurred images are presented. The second task was a newly invented paradigm designed to investigate neural and behavioral responses to images depicting social situations. For this purpose, participants viewed images of various social and non-social scenes with varying numbers of people. After each image, participants were instructed to indicate how much they would like to be in the depicted situation.

Functional connectivity analyses were performed on the MID/SID task. This task is known to activate the dopaminergic reward system which plays an important role in both AUD and stress. As in previous studies (13,14), all task-related activity was removed by regressing out hemodynamic responses induced by the task. Regressors for the task included the onset times for each of the four conditions (monetary and social conditions both with and without reward) as well as a fifth regressor containing the onset times of the received feedback for all trials. Additionally, the GLM included the six realignment parameters, their temporal derivatives, and a scrubbing regressor, removing all volumes with a framewise displacement > 1 mm.

This approach must be taken into account when interpreting the results. However, it may also offer advantages over classic resting-state studies: As functional connectivity depends on the current state of a person and the state is not well defined during rest (15), individual differences might emerge due to differences in spontaneous mental activity. In contrast, in the design of the present study, the investigation takes place in a controlled state of reward processing.

**fMRI: Seed masks**

Masks for the dorsal striatum, thalamus, and amygdala were extracted from the Harvard-Oxford Atlas (HOA, (16,17)). Masks of the insula and ACC were taken from the Brainnetome Atlas (18) as they are smaller and better fitted the desired regions of interest than those in the HOA in the normalized images. Masks for the anterior and posterior DMN were obtained from Smallwood et al. (19). Following previous studies of our own and other research groups (20–25), masks of the ventral striatum were defined as spheres with an 8 mm radius around peak coordinates (Montreal Neurological Institute coordinates: left, −10, 10, −2; right, 12, 14, −4) from a meta-analysis examining reward anticipation in the ventral striatum ((20–24)). Except for the ACC and DMN, all analyses were performed for the left and right hemispheres individually.

### **Results: Functional connectivity differences between groups on the control day**

Table S1. Group differences on control day only (controls > patients)

|  | side | MNI coordinates | | | Cluster size | T | p (peak) |
| --- | --- | --- | --- | --- | --- | --- | --- |
|  |  | x | y | z |  |  |  |
| **Seed: left ventral striatum** | | | | | | | |
| angular gyrus, middle temporal gyrus, inferior + superior parietal lobule, supramarginal gyrus | L | -51 | -58 | 26 | 290 | 5.61 | 0.001 |
|  |  | -57 | -52 | 23 |  | 5.29 |  |
|  |  | -39 | -55 | 44 |  | 4.65 |  |
| precuneus, midcingulate cortex | L/R | 0 | -43 | 56 | 402 | 5.18 | <0.001 |
|  |  | 0 | -34 | 62 |  | 4.82 |  |
|  |  | 6 | -46 | 65 |  | 4.61 |  |
| **Seed: right ventral striatum** |  |  |  |  |  |  |  |
| precuneus, midcingulate cortex | L | 0 | -43 | 56 | 184 | 4.53 | 0.011 |
|  |  | -6 | -43 | 44 |  | 3.78 |  |
|  |  | 0 | -34 | 35 |  | 3.67 |  |
| superior + middle frontal gyrus, superior medial gyrus | L/R | -12 | 50 | 35 | 178 | 4.42 | 0.011 |
|  |  | -24 | 44 | 32 |  | 4.23 |  |
|  |  | 3 | 47 | 35 |  | 3.71 |  |
| **Seed: left dorsal striatum** | | | | | | | |
| angular gyrus, superior + inferior parietal lobule, midcingulate cortex | L/R | 36 | -58 | 47 | 3395 | 5.87 | <0.001 |
|  |  | -48 | -61 | 29 |  | 5.76 |  |
|  |  | -39 | -55 | 44 |  | 5.48 |  |
| middle temporal gyrus | R | 66 | -40 | 5 | 148 | 5.75 | 0.020 |
|  |  | 60 | -49 | -7 |  | 4.51 |  |
|  |  | 27 | -70 | -28 | 1860 | 5.56 | <0.001 |
| cerebellum, lingual gyrus | L/R | -30 | -76 | -31 |  | 5.24 |  |
|  |  | 18 | -76 | -25 |  | 5.09 |  |
| superior frontal gyrus | L | -15 | 50 | 35 | 128 | 4.39 | 0.026 |
|  |  | -18 | 41 | 35 |  | 4.01 |  |
| superior medial gyrus, superior frontal gyrus | R | 9 | 62 | 23 | 124 | 4.02 | 0.026 |
|  |  | 0 | 44 | 29 |  | 4.01 |  |
|  |  | 24 | 41 | 38 |  | 3.68 |  |
| **Seed: right dorsal striatum** |  |  |  |  |  |  |  |
| cerebellum | R | 30 | -67 | -28 | 270 | 4.86 | 0.002 |
|  |  | 39 | -73 | -31 |  | 4.35 |  |
|  |  | 18 | -76 | -25 |  | 4.30 |  |
| superior + inferior parietal lobule, angular gyrus | L | -21 | -73 | 53 | 240 | 4.77 | 0.002 |
|  |  | -42 | -52 | 53 |  | 4.37 |  |
|  |  | -51 | -61 | 29 |  | 4.12 |  |
| superior parietal lobule, angular gyrus, precuneus | R | 33 | -61 | 53 | 124 | 4.54 | 0.018 |
|  |  | 39 | -58 | 47 |  | 4.17 |  |
|  |  | 9 | -58 | 53 |  | 3.94 |  |
| superior parietal lobule, postcentral gyrus | R | 42 | -43 | 56 | 121 | 4.27 | 0.018 |
|  |  | 30 | -46 | 62 |  | 4.04 |  |
|  |  | 24 | -40 | 62 |  | 3.72 |  |
| precuneus, postcentral gyrus, superior parietal lobule | L | -15 | -46 | 71 | 102 | 4.14 | 0.022 |
|  |  | -36 | -34 | 59 |  | 3.70 |  |
|  |  | -24 | -43 | 65 |  | 3.62 |  |
| precuneus, midcingulate cortex | L/R | 0 | -43 | 56 | 132 | 4.12 | 0.018 |
|  |  | 3 | -40 | 47 |  | 3.92 |  |
|  |  | 3 | -46 | 32 |  | 3.33 |  |
| lingual gyrus | L/R | -9 | -40 | -1 | 105 | 4.11 | 0.022 |
|  |  | 18 | -28 | -10 |  | 4.08 |  |
|  |  | -12 | -55 | 2 |  | 3.98 |  |
| cerebellum | L | -27 | -76 | -28 | 120 | 4.08 | 0.018 |
|  |  | -15 | -70 | -19 |  | 4.05 |  |
| superior frontal gyrus | L | -21 | 38 | 35 | 109 | 4.07 | 0.022 |
|  |  | -12 | 50 | 35 |  | 4.04 |  |
| **Seed: right dorsal insula** | | | | | | | |
| cerebellum | R | 18 | -76 | -25 | 145 | 4.36 | 0.047 |
|  |  | 42 | -58 | -43 |  | 4.12 |  |
|  |  | 30 | -67 | -28 |  | 3.56 |  |
| **Seed: anterior DMN** | | | | | | |  |
| putamen, caudate nucleus, superior temporal gyrus, pallidum | L/R | -15 | 11 | 2 | 14228 | 7.43 | <0.001 |
|  |  | 15 | 8 | 8 |  | 6.61 |  |
|  |  | 30 | -4 | 8 |  | 6.58 |  |
| **Seed: posterior DMN** | | | | | | | |
| putamen, inferior parietal lobule, caudate nucleus | L | -24 | 8 | 2 | 1612 | 5.51 | <0.001 |
|  |  | -42 | -25 | 44 |  | 5.11 |  |
|  |  | -27 | -19 | 2 |  | 5.04 |  |
| caudate nucelus, pallidum | R | 9 | 5 | 8 | 195 | 5.27 | 0.003 |
|  |  | 21 | 5 | -1 |  | 4.58 |  |
| precentral gyrus, inferior frontal gyrus | R | 45 | -1 | 44 | 131 | 4.64 | 0.011 |
|  |  | 51 | 8 | 32 |  | 4.10 |  |
|  |  | 54 | -4 | 38 |  | 3.84 |  |
| postcentral gyrus, supramarginal gyrus, inferior parietal lobule | R | 39 | -31 | 47 | 177 | 4.38 | 0.004 |
|  |  | 60 | -34 | 44 |  | 4.20 |  |
|  |  | 48 | -22 | 44 |  | 3.35 |  |
| rolandic operculum, insula lobe, supramarginal gyrus | R | 42 | -28 | 20 | 85 | 4.25 | 0.036 |
|  |  | 63 | -19 | 29 |  | 3.79 |  |
|  |  | 57 | -25 | 23 |  | 3.78 |  |
| precuneus, midcingulate cortex, paracentral lobule | L/R | -3 | -40 | 56 | 91 | 4.10 | 0.035 |
|  |  | 3 | -37 | 47 |  | 4.05 |  |
|  |  | -3 | -25 | 50 |  | 3.84 |  |
| midcingulate cortex, posterior-medial frontal | R | 0 | -7 | 47 | 78 | 3.86 | 0.041 |
|  |  | 6 | -10 | 62 |  | 3.70 |  |

p < 0.001 (unc.), FDR, cluster-level, DMN = Default Mode Network

### **References**

1. Saunders JB, Aasland OG, Babor TF, De La Fuente JR, Grant M (06/1993): Development of the Alcohol Use Disorders Identification Test (AUDIT): WHO Collaborative Project on Early Detection of Persons with Harmful Alcohol Consumption-II. *Addiction* 88: 791–804.

2. Kirschbaum C, Pirke KM, Hellhammer DH (1993): The “Trier Social Stress Test”--a tool for investigating psychobiological stress responses in a laboratory setting. *Neuropsychobiology* 28: 76–81.

3. Allen AP, Kennedy PJ, Dockray S, Cryan JF, Dinan TG, Clarke G (2017): The Trier Social Stress Test: Principles and practice. *Neurobiol Stress* 6: 113–126.

4. Het S, Rohleder N, Schoofs D, Kirschbaum C, Wolf OT (8/2009): Neuroendocrine and psychometric evaluation of a placebo version of the “Trier Social Stress Test.” *Psychoneuroendocrinology* 34: 1075–1086.

5. Grimm J (2009): *State-Trait-Anxiety Inventory Nach Spielberger. Deutsche Lang-Und Kurzversion. Methodenforum Der Universität Wien*. MF-Working Paper 2009/02.

6. Hautzinger M, Keller F, Kühner C (2006): *Beck Depressions-Inventar (BDI-II)*. Harcourt Test Services.

7. Beck AT, Steer RA, Brown GK (2013): Beck Depressions-Inventar-FS. *Deutsche Bearbeitung von Sören Kliem & Elmar*.

8. Schulz P, Jansen LJ, Schlotz W (2005): Stressreaktivität: Theoretisches Konzept und Messung. *Diagnostica* 51: 124–133.

[9. Schulz P, Schlotz W, Becker P, Schulz P, Schlotz W, Becker P (2004): Trierer Inventar zum Chronischen Stress (TICS) [Trier Inventory for Chronic Stress (TICS)]. Retrieved February 15, 2023, from](http://paperpile.com/b/sZ4Wbk/daVao) <https://eprints.soton.ac.uk/50017/>

10. Knutson B, Westdorp A, Kaiser E, Hommer D (2000): FMRI visualization of brain activity during a monetary incentive delay task. *Neuroimage* 12: 20–27.

11. Rademacher L, Krach S, Kohls G, Irmak A, Gründer G, Spreckelmeyer KN (2010): Dissociation of neural networks for anticipation and consumption of monetary and social rewards. *Neuroimage* 49: 3276–3285.

12. Spreckelmeyer KN, Krach S, Kohls G, Rademacher L, Irmak A, Konrad K, *et al.* (2009): Anticipation of monetary and social reward differently activates mesolimbic brain structures in men and women. *Soc Cogn Affect Neurosci* 4: 158–165.

13. Esslinger C, Walter H, Kirsch P, Erk S, Schnell K, Arnold C, *et al.* (2009): Neural Mechanisms of a Genome-Wide Supported Psychosis Variant. *Science* 324: 605.

14. Paulus FM, Krach S, Bedenbender J, Pyka M, Sommer J, Krug A, *et al.* (2013): Partial support for ZNF804A genotype-dependent alterations in prefrontal connectivity. *Hum Brain Mapp* 34: 304–313.

15. Meer JN van der, Breakspear M, Chang LJ, Sonkusare S, Cocchi L (2020): Movie viewing elicits rich and reliable brain state dynamics. *Nat Commun* 11: 5004.

16. Desikan RS, Ségonne F, Fischl B, Quinn BT, Dickerson BC, Blacker D, *et al.* (2006): An automated labeling system for subdividing the human cerebral cortex on MRI scans into gyral based regions of interest. *Neuroimage* 31: 968–980.

17. Frazier JA, Chiu S, Breeze JL, Makris N, Lange N, Kennedy DN, *et al.* (2005): Structural brain magnetic resonance imaging of limbic and thalamic volumes in pediatric bipolar disorder. *Am J Psychiatry* 162: 1256–1265.

18. Fan L, Li H, Zhuo J, Zhang Y, Wang J, Chen L, *et al.* (2016): The Human Brainnetome Atlas: A New Brain Atlas Based on Connectional Architecture. *Cereb Cortex* 26: 3508–3526.

19. Smallwood J, Bernhardt BC, Leech R, Bzdok D, Jefferies E, Margulies DS (2021): The default mode network in cognition: a topographical perspective. *Nat Rev Neurosci* 22: 503–513.

20. Seitz KI, Ueltzhöffer K, Rademacher L, Paulus FM, Schmitz M, Herpertz SC, Bertsch K (2023): Your smile won’t affect me: Association between childhood maternal antipathy and adult neural reward function in a transdiagnostic sample. *Transl Psychiatry* 13: 70.

21. Diekhof EK, Kaps L, Falkai P, Gruber O (2012): The role of the human ventral striatum and the medial orbitofrontal cortex in the representation of reward magnitude - an activation likelihood estimation meta-analysis of neuroimaging studies of passive reward expectancy and outcome processing. *Neuropsychologia* 50: 1252–1266.

22. Eckstrand KL, Forbes EE, Bertocci MA, Chase HW, Greenberg T, Lockovich J, *et al.* (2019): Anhedonia Reduction and the Association Between Left Ventral Striatal Reward Response and 6-Month Improvement in Life Satisfaction Among Young Adults. *JAMA Psychiatry* 76: 958–965.

23. Chase HW, Nusslock R, Almeida JR, Forbes EE, LaBarbara EJ, Phillips ML (2013): Dissociable patterns of abnormal frontal cortical activation during anticipation of an uncertain reward or loss in bipolar versus major depression. *Bipolar Disord* 15: 839–854.

24. Chase HW, Fournier JC, Bertocci MA, Greenberg T, Aslam H, Stiffler R, *et al.* (2017): A pathway linking reward circuitry, impulsive sensation-seeking and risky decision-making in young adults: identifying neural markers for new interventions. *Transl Psychiatry* 7: e1096.

25. Mayer AV, Preckel K, Ihle K, Piecha FA, Junghanns K, Reiche S, *et al.* (2022): Assessment of Reward-Related Brain Function After a Single Dose of Oxytocin in Autism: A Randomized Controlled Trial. *Biol Psychiatry Glob Open Sci* 2: 136–146.
